# *MONET*: a toolbox integrating top-performing methods for network modularisation

**DOI:** 10.1101/611418

**Authors:** Mattia Tomasoni, Sergio Gómez, Jake Crawford, Weijia Zhang, Sarvenaz Choobdar, Daniel Marbach, Sven Bergmann

## Abstract

**Summary:** We define a disease module as a partition of a molecular network whose components are jointly associated with one or several diseases or risk factors thereof. Identification of such modules, across different types of networks, has great potential for elucidating disease mechanisms and establishing new powerful bio-markers. To this end, we launched the “Disease Module Identification (DMI) DREAM Challenge”, a community effort to build and evaluate unsupervised molecular network modularisation algorithms (Choobdar *et al*., 2018). Here we present *MONET*, a toolbox providing easy and unified access to the three top methods from the DMI DREAM Challenge for the bioinformatics community.

**Availability and Implementation:** *MONET* is a command line tool for Linux, based on Docker and Singularity containers; the core algorithms were written in R, Python, Ada and C++. It is freely available for download at https://github.com/BergmannLab/MONET.git

**Contact:** (MT); (SB)

## 1 Introduction

Gene networks, such as protein interaction, signalling, gene co-expression and homology networks, provide scaffolds of linked genes. Sub-networks, or modules, include genes normally acting in concert but whose joint function may be disrupted, if any of its members is missing, or dis-regulated. For Disease Modules this disruption can lead to a disease phenotype. The identification of such modules is therefore useful for elucidating disease mechanisms and establishing new bio-markers and potential therapeutic targets. Yet, which methods work best to extract such modules from different types of networks is not well understood. This prompted us to initiate the “Disease Module Identification (DMI) DREAM Challenge” (Choobdar *et al*., 2018), providing an unbiased and critical assessment of 75 contributed module identification methods. Our method evaluation used summary statistics from more than 200 disease relevant Genome-wide Association Studies (GWAS) in conjunction with our Pascal tool (Lamparter *et al*., 2016), avoiding the bias of using annotated molecular pathways.

The top-performing methods implemented novel algorithms that advanced the state of the art, clearly outperforming off-the-shelf tools. We therefore decided to make the top three methods available for the bioinformatics community in a single user-friendly package: *MONET* is a command line tool based on *Docker* and *Singularity* virtualization technologies, automatically installing the tool with all its dependencies inside a *container*, avoiding time-consuming and error-prone manual installations of computing environments and libraries. All computations then take place in this *sandbox* environment and once the output is ready, all resources can be fully released bringing the user’s machine back to its original state.

## 2 Methods and implementation

While our challenge was able to establish *Kernel Clustering Optimization using the “Diffusion State Distance” metric* by Cao *et al*. (2014) (hereafter K1) as the overall winner, there were several strong competitors using entirely different approaches for the network modularisation. Importantly, we observed that no single method was superior on all network types and that Disease Modules identified by different methods were often complementary (Choobdar *et al*., 2018).

### 2.1 K1: Top method using kernel clustering

K1 is based on the “Diffusion State Distance” (DSD), a novel graph metric which is built on the premise that paths through low-degree nodes are stronger indications of functional similarity than paths that traverse high-degree nodes by Cao *et al*. (2014). The DSD metric is used to define a pairwise distance matrix between all nodes, on which a spectral clustering algorithm is applied. In parallel, dense bipartite sub-graphs are identified using standard graph techniques. Finally, results are merged into a single set of non-overlapping clusters.

BLOG: https://www.synapse.org/#!Synapse:syn7349492/wiki/407359

### 2.2 M1: Top method using modularity optimization

M1 employs an original technique named *Multiresolution* introduced by (Arenas *et al*., 2008) to explore all topological scales at which modules may be found. The novelty of this approach relies on the introduction of a parameter, called *resistance*, which controls the aversion of nodes to form modules. Modularity (Newman and Girvan, 2004; Arenas *et al*., 2007) is optimized using an ensemble of algorithms: Extremal optimization (Duch and Arenas, 2005), Spectral optimization (Newman, 2006), Fast algorithm (Newman, 2004), Tabu search (Arenas *et al*., 2008), and fine-tuning by iterative repositioning of individual nodes in adjacent modules.

BLOG: https://www.synapse.org/#!Synapse:syn7352969/wiki/407384

### 2.3 R1: Top method using random walk

R1 is based on a variant of *Markov Cluster Algorithm* known as balanced Multi-layer Regularized Markov Cluster Algorithm (*bMLRMCL*) (Satuluri *et al*., 2010) which scales well to large graphs and minimizes the number of oversized clusters. First, a pre-processing step is applied so that edges with low weights are discarded and all remaining edges are scaled to integer values. Then, *bMLRMCL* is applied iteratively on modules of size grater than 100 nodes.

BLOG: https://www.synapse.org/#!Synapse:syn7286597/wiki/406659

## 3 Performance

Figure 1 illustrates the performance of the *MONET* algorithms on simulated graphs with planted community structure, generated using the class of benchmark graphs proposed by Lancichinetti *et al*. (2008). Modularisation performance is measured using Normalized Mutual Information (NMI). Experiments were carried out on regular desktop hardware. In accordance with performance evaluations within the DMI DREAM Challenge, K1, the winner, requires the most computational resources, with a runtime of about one day and the highest memory allocation for processing on the Challenge inputs. M1, the second runner-up, completed the Challenge in a few hours and displayed excellent performance on the simulated benchmark (even superior to K1, especially in case of extremely high fraction of inter-module edges and extremely low memory requirements). R1, the second runner-up, is the only method that requires parameters to be tuned (nine in total); nevertheless, we believe it is an excellent addition to our tool, as it performed close to K1/M1 on the benchmark, it requires only moderate memory and has an extremely low run time (it completed the Challenge in under an hour).

**Fig. 1.**
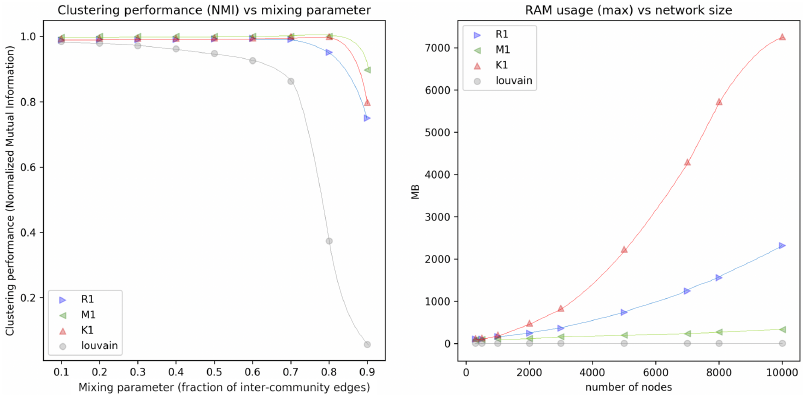
Comparison of the MONET methods (K1, M1 and R1) against a baseline (Louvain) on simulated graphs with planted community structure. On the left: clustering performance (NMI) as a function of the fraction of inter-module edges (mixing parameter). Right: memory requirements as a function of network size. Each point represents an average of the results obtained performing a grid search over the following parameter space (at least two repetitions for each combination of parameters): number of nodes: 5k, 7k, 8k, 10k; average node degree: 15, 20, 25; exponent of the distribution of community sizes: 1, 2; exponent of the distribution of node degrees: 2, 3.

## 4 Installation and usage

*MONET* is extremely simple to install/uninstall and run. The only requirement is having installed either Docker (Merkel, 2014) or Singularity (Kurtzer *et al*., 2017). For detailed instructions and information about usage and I/O formats, please refer to the README file on the github repository.

~~~
$ git clone https://github.com/BergmannLab/MONET.git
$ cd MONET && ./install.sh
$ monet --help
$ monet --method=M1 --container=docker \
 --input=./input/network.txt --output=./output
~~~

## Notes

#### Summary of Updates

Changing the title.

https://www.synapse.org/#!Synapse:syn7543444/wiki/409344

